# Insights into Cross-species Evolution of Novel Human Coronavirus SARS-CoV-2 and Defining Immune Determinants for Vaccine Development

**DOI:** 10.1101/2020.01.29.925867

**Authors:** Arunachalam Ramaiah, Vaithilingaraja Arumugaswami

## Abstract

Novel severe acute respiratory syndrome coronavirus 2 (SARS-CoV-2) outbreak in the city of Wuhan, China during December 2019, has now spread to various countries across the globe triggering a heightened containment effort. This human pathogen is a member of betacoronavirus genus carrying 30 kilobase of single positive-sense RNA genome. Understanding the evolution, zoonotic transmission, and source of this novel virus would help accelerating containment and prevention efforts. The present study reported detailed analysis of SARS-CoV-2 genome evolution and potential candidate peptides for vaccine development. This new coronavirus genotype might have been evolved from a bat-coronavirus by accumulating non-synonymous mutations, indels, and recombination events. Structural proteins Spike (S), and Membrane (M) had extensive mutational changes, whereas Envelope (E) and Nucleocapsid (N) proteins were very conserved suggesting differential selection pressures exerted on SARS-CoV-2 during evolution. Interestingly, SARS-CoV-2 Spike protein contains a 39 nucleotide sequence insertion relative to SARS-like bat-SL-CoVZC45/2017. Furthermore, we identified eight high binding affinity (HBA) CD4 T-cell epitopes in the S, E, M and N proteins, which can be commonly recognized by HLA-DR alleles of Asia and Asia-Pacific Region population. These immunodominant epitopes can be incorporated in universal subunit SARS-CoV-2 vaccine. Diverse HLA types and variations in the epitope binding affinity may contribute to the wide range of immunopathological outcomes of circulating virus in humans. Our findings emphasize the requirement for continuous surveillance of SARS-CoV-2 strains in live animal markets to better understand the viral adaptation to human host and to develop practical solutions to prevent the emergence of novel pathogenic SARS-CoV-2 strains.

## INTRODUCTION

Current severe acute respiratory syndrome coronavirus 2 (SARS-CoV-2) outbreak in China and its subsequent spread to multiple countries across the continents precipitated a major global health crisis. Uncovering the SARS-CoV-2 genome evolution and immune determinants of its proteome would be crucial for outbreak containment and preventative efforts. Coronaviruses (CoVs) are single positive-stranded RNA viruses belonging to *Coronaviridae* family^1^, which are divided into four genera: *Alpha-, Beta-, Delta- and Gamma-coronavirus*. Coronaviruses are commonly found in many species of animals, including bats, camels and humans. Occasionally, the animal coronaviruses can acquire genetic mutations by errors during genome replication or recombination mechanism, which can further expand their tropism to humans. The first human coronavirus was discovered in the mid-1960s. A total of six human coronavirus types were identified to be responsible for causing human respiratory ailments, which include two alpha coronaviruses (229E; NL63), and four beta CoVs (OC43; HKU1; SARS-CoV; MERS-CoV)^2^. Typically, these coronaviruses cause asymptomatic infection or severe acute respiratory illness, including fever, cough, and shortness of breath. However, other symptoms such as gastroenteritis and neurological diseases of varying severity have also been reported^3–5^.

The recent two coronavirus human outbreaks were caused by betacoronaviruses, SARS-CoV (Severe Acute Respiratory Syndrome-CoV) and MERS-CoV (Middle East Respiratory Syndrome-CoV). In 2002, SARS-CoV outbreak was first reported in China, which spread globally causing hundreds of deaths with a 11% mortality rate. In 2012, MERS-CoV was first recognized in Saudi Arabia and subsequently in other countries, which has a fatality rate of 37%. Since December 2019, the seventh novel human betacoronavirus SARS-CoV-2 causing pneumonia outbreak was reported by the World Health Organization. This novel virus is linked with an outbreak of febrile respiratory illness in the city of Wuhan, Hubei Province, China with an epidemiological association to the Huanan Seafood Wholesale Market selling live farm and wild animals^6^ and elsewhere in the city^1^.

While the SARS-CoV-2 appears to be a zoonotic infectious disease, the specific animal species and other reservoirs need to be clearly defined. Identification of the source animal species for this outbreak would facilitate global public health authorities to inspect the trading route and the movement of wild and domestic animals to Wuhan and taking control measures to limit the spread of this disease^7^. The SARS-CoV-2 outbreak has a fatality rate of 2% with significant impact on global health and economy. As of 29 January 2020, the SARS-CoV-2 has spread to fifteen countries including Asia-Pacific region (Thailand, Japan, The Republic of Korea, Malaysia, Cambodia, Sri Lanka, Nepal, Vietnam, Singapore, and Australia), United Arab Emirates, France, Germany, Canada and the United States with 6065 confirmed cases, in which 5997 reported from China^8^.

Here, we report a detailed genetic analysis of SARS-CoV-2 evolution based on 47 genome sequences available in Global Initiative on Sharing All Influenza Database (GISAID) (https://www.gisaid.org/) (Supplemental Table 1) and one genome sequence from GenBank. We further defined potential epitope targets in the structural proteins (SPs) for vaccine design.

## MATERIALS AND METHODS

### Phylogenetic tree construction

We employed the following methodologies for genetic analyses. The genome sequences of SARS-CoV-2 were obtained on 29 January 2020 along with genetically related 10 SARS-CoV/SARS-CoV-like virus genome sequences that we identified through the BLASTN search in NCBI database. We also included 10 sequences of MERS-CoV strains and one outgroup alphacoronavirus sequence. SARS-CoV-2 genome (30 Kb in size) codes for a large non-structural polyprotein ORF1ab that is proteolytically cleaved to generate 15/16 proteins, and four structural proteins - Spike (S), Envelope (E), Membrane (M) and Nucleocapsid (N), as well as five accessory proteins (ORF3a, ORF6, ORF7a, ORF8 and ORF10). The genome sequences were aligned using the MAFFTv7 program^9^, which were exported to MEGA-X^10^ for Neighbor-Joining (NJ) phylogenetic tree construction with 1000 bootstrap support and genetic distance calculations. Structural genes/proteins were aligned with closely related sequences using Clustal Omega Program^11^.

### High Binding Affinity (HBA) epitope identification

In order to identify the potential target peptides for vaccine design, the structural protein sequences of highly conserved and annotated representative SARS-CoV-2 strain Wuhan-Hu-1 (MN908947.3) were used to predict the CD4 T-cell epitopes (TCEs) with the Immune Epitope Database and Analysis Resource (IEDB) consensus tool by IEDB recommended prediction method^12,13^. To identify the most immunodominant (IMD) CD4 epitopes, we employed the computational approach described previously^14^. For all the possible 15-mer peptides from the SPs, the binding affinity was predicted using six most prevalent Human Leukocyte Antigen (HLA; MHC Class-II DR) alleles (http://allelefrequencies.net/) that cover most of the ethnic population in China (PRC), Thailand (TH) and Japan (JPN) (Supplemental Table 2). Since, the prevalence of dominant HLA-DR types may vary in human populations from different countries, the prevalent six HLA-DR alleles covering Asia-Pacific region^14^ and Global population data sets have been used for binding affinity predictions to assess the generality of those predicted epitopes with high binding affinities (HBA). We calculated the average binding affinity score for each predicted 15-mer peptides against all the predominant HLA-DR types from each country/region using a sliding window approach (Figure 2). This approach is more efficient as the epitopes predicted with average HBA (IC <=10 nM), indicate that those epitopes are more likely to bind and enhance a cellular immune response in the studied population.

**Figure 1.**
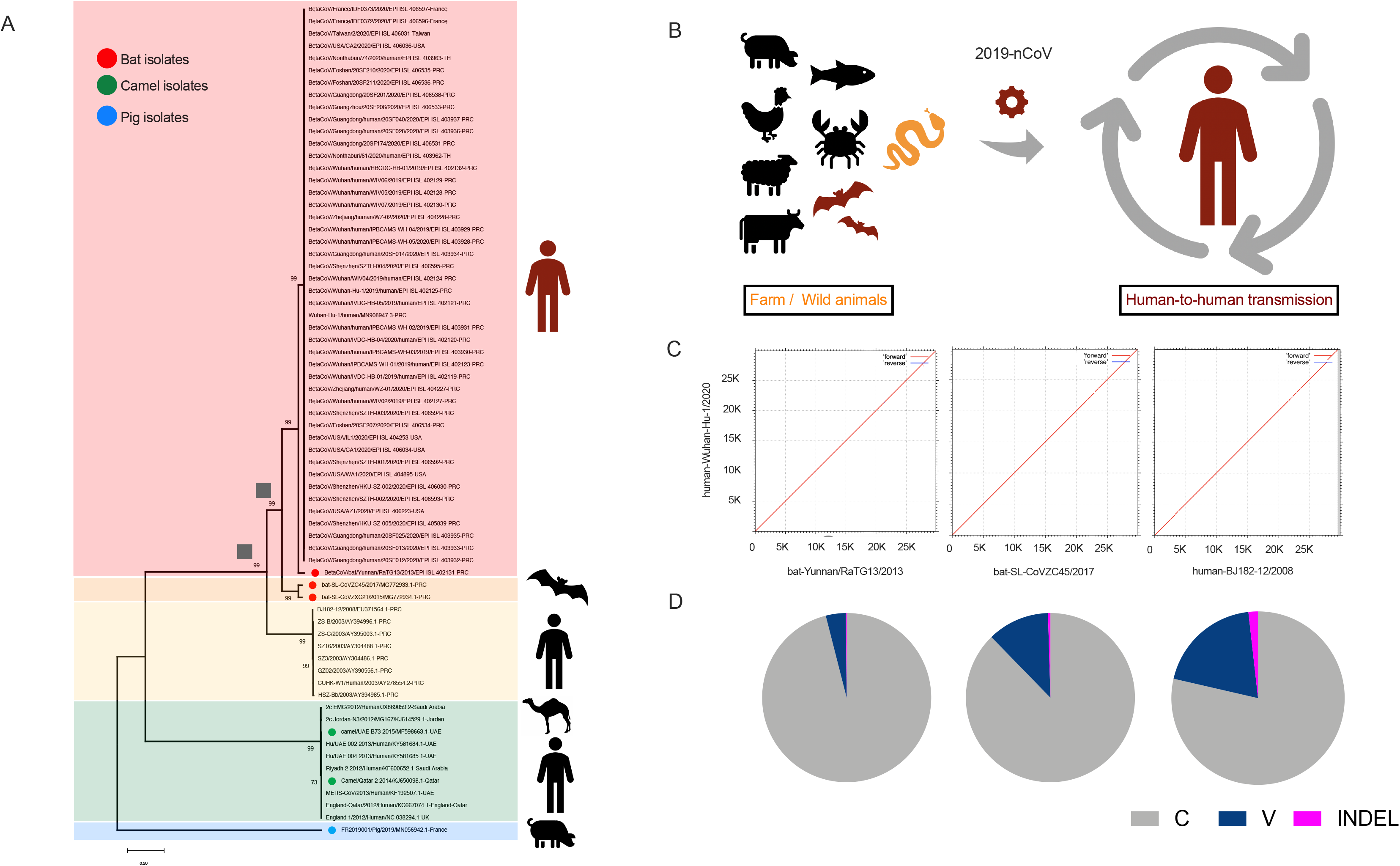
Phylogenetic relationships of novel SARS-CoV-2 (*2019-nCoV*) viral strains and the possible route of viral evolution and disease transmission. (A) NJ bootstrap consensus tree was reconstructed based on the genome sequences with 1000 bootstrap replications. SARS-CoV-2 strains (n=48 including one bat strain) from current outbreak, SARS-like-CoV (n=2), SARS-CoV (n=8) and MERS-CoV (n=10) viruses are colored in red, orange, yellow and green, respectively. The alphacoronavirus (MN056942) was used as an outgroup control (blue). (B) Evolution of a SARS-CoV-2 virus from bat/unknown animals to humans and spread among humans. (C) Dot plots comparison between the human SARS-CoV-2 (MN908947.3) and bat/Yunnan/RaTG13/2013 (EPI-ISL-402131) (left), or bat SARS-CoV-like (MG772933) viruses (middle); or human SARS-CoV (EU371564) (right). All these viruses were reported from China. (D) The proportion of conserved, variable (Single Nucleotide Polymorphism) and INDELs in SARS-CoV-2 was uncovered by aligning it to bat-coronavirus (left), bat SARS-CoV-like (middle) and human SARS-CoV (right).

**Figure 2.**
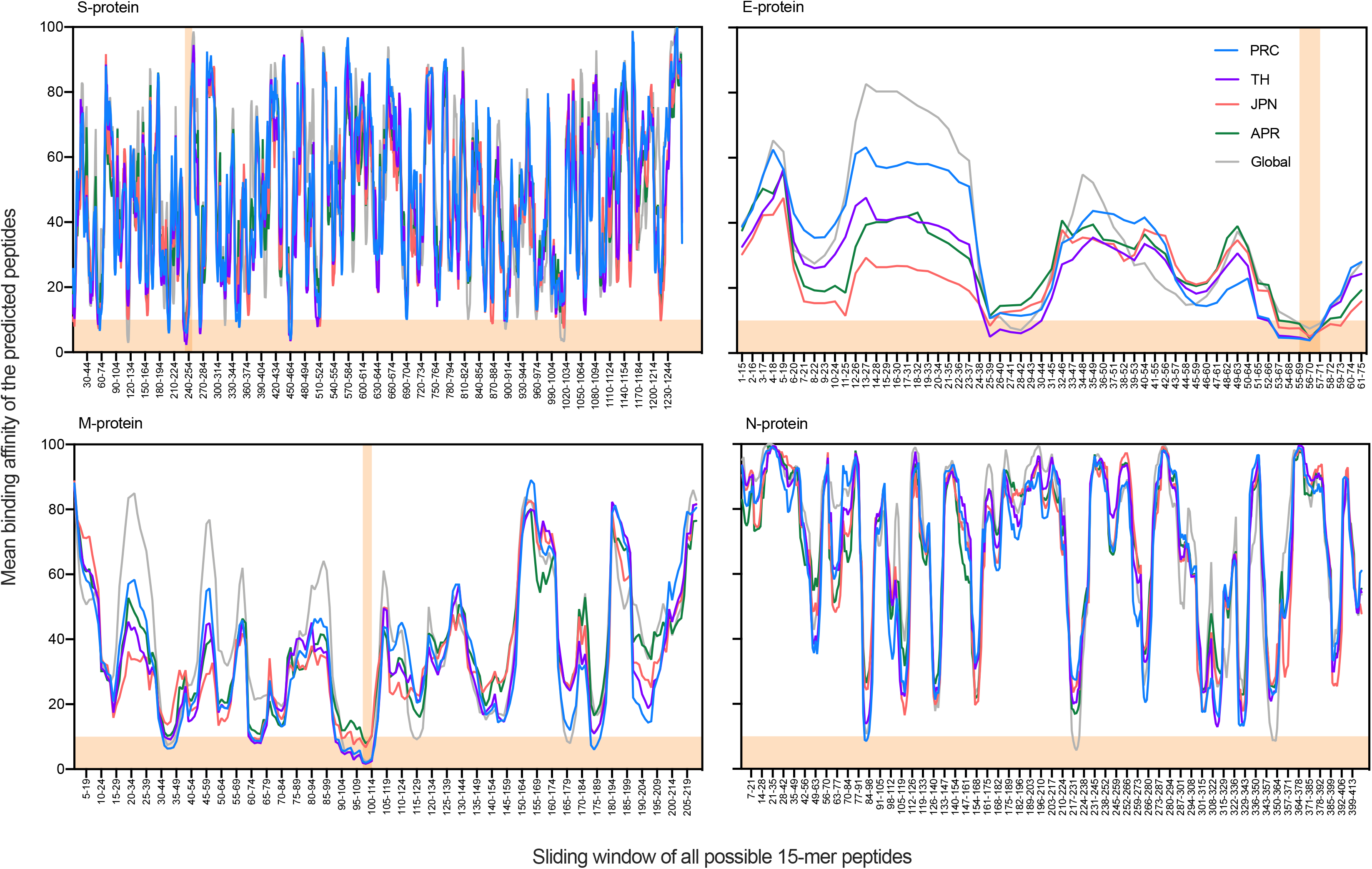
Comparison of SARS-CoV-2 structural proteins’ epitopes with mean HBA to predominant HLA-DR alleles from China (PRC), Thailand (TH), Japan (JPN), Asia-Pacific region (APR) and Global population. The mean binding affinity score was calculated from the binding affinity (score) of each predicted 15-mer HBA peptides (sliding window approach) from four structural proteins with six predominant HLA-DR types from China (PRC), Thailand (TH), Japan (JPN), Asia-Pacific region (APR) and Global population sets. Functional islands with mean HBA TCEs (IC <= 10nM) in the structural proteins were located in the horizontal orange color. The epitopes having mean HBA to all five population HLA-DR alleles are highlighted with vertical orange bars.

## RESULTS AND DISCUSSION

### Phylogenetic analysis reveals SARS-CoV-2 closely related to Bat coronavirus

Genome based phylogenetic tree analysis revealed a major cluster of large clade was formed with humans SARS-CoV-2 strains, SARS-CoV-like strains from bats and human SARS-CoV viruses (Figure 1A) with 100 bootstrap support. All SARS-CoV-2 strains were clustered together with 96% bootstrap support but separated from bat/bat-SARS like strains and human SARS-CoVs clusters, indicating that SARS-CoV-2 belongs to a novel genotype. A small cluster was formed by MERS-CoV strains, which was genetically unrelated to the strains circulating in the current outbreak. This result indicated that the causative agent of current outbreak is a betacoronavirus that is genetically and evolutionarily related to bat coronaviruses: bat/Yunnan/RaTG13/2013 and bat/SARS-CoV-like-viruses. Moreover, the result suggests that the evolutionary pressures exerted on SARS-CoV-2, as well as bat and human SARS-CoVs were different. Interestingly, strains from the current outbreak were clearly segregated from bat SARS-CoV-like viruses (isolated in 2015 and 2017), but located adjacent to the bat/Yunnan/RaTG13/2013 virus isolated in 2013. Besides, both SARS-CoV-2 and bat coronavirus clusters share a common ancestor (square), which is supported by 92% bootstrap replications in the phylogeny. The ancestors of both human SARS-CoV and SARS-CoV-2 share a common ancestor. Overall our data indicates that the SARS-CoV-2 is not closely clustering with either human SARS-CoV or bat-SARS-CoV like strains but shares a common ancestor.

The mean group genetic distance among SARS-CoV-2 strains was zero, indicating high genetic conservation among currently circulating viruses. Similar findings were observed in human SARS and MERS isolates, but bat SARS-like isolates showed a genetic distance of 0.04. The current circulating strains have close genetic associations with the 2013 bat coronavirus, bat/Yunnan/RaTG13/2013 (EPI-ISL-402131). The SARS-CoV-2 Wuhan-Hu-1 and bat/Yunnan/RaTG13/2013 strains share 96% sequence identity (genetic distance 0.04) with 1141 variable nucleotide substitutions and 48 INDELs (Figure 1C and D; Table 1A). While, bat-SL-CoVZC45/2017 virus and SARS human-BJ182-12/2008 virus from other clusters had 88% and 80% sequence identity (distance 0.23 and 0.46), 3542 and 5879 nucleotide substitutions and 137 and 535 INDELs, respectively with SARS-CoV-2 Wuhan-Hu-1, suggesting that bat/Yunnan/RaTG13/2013 strain is likely the most recent ancestor of currently circulating SARS-CoV-2 strains. This most recent ancestral coronavirus strain from bat species underwent significant evolution during 2013 to 2019 possibly by genetic shift before spreading to humans (Figure 1B). However, it is likely that coronaviruses from other animal species may also be a source for this outbreak. Comparison of Spike protein of SARS-CoV-2 and bat coronavirus bat/Yunnan/RaTG13/2013 showed 97.7% sequence identity with 29 non-synonymous changes and 4 amino acid insertions (position 681-684, PRRA) (Table 1C). Whereas, M protein shared 98.6% identity with one amino acid insertion (position 4, S) and N protein showed 99% identity with 4 non-synonymous changes (P37S, S215G, S243G, Q267A). Envelope protein shares 100% identity.

**Table 1.**
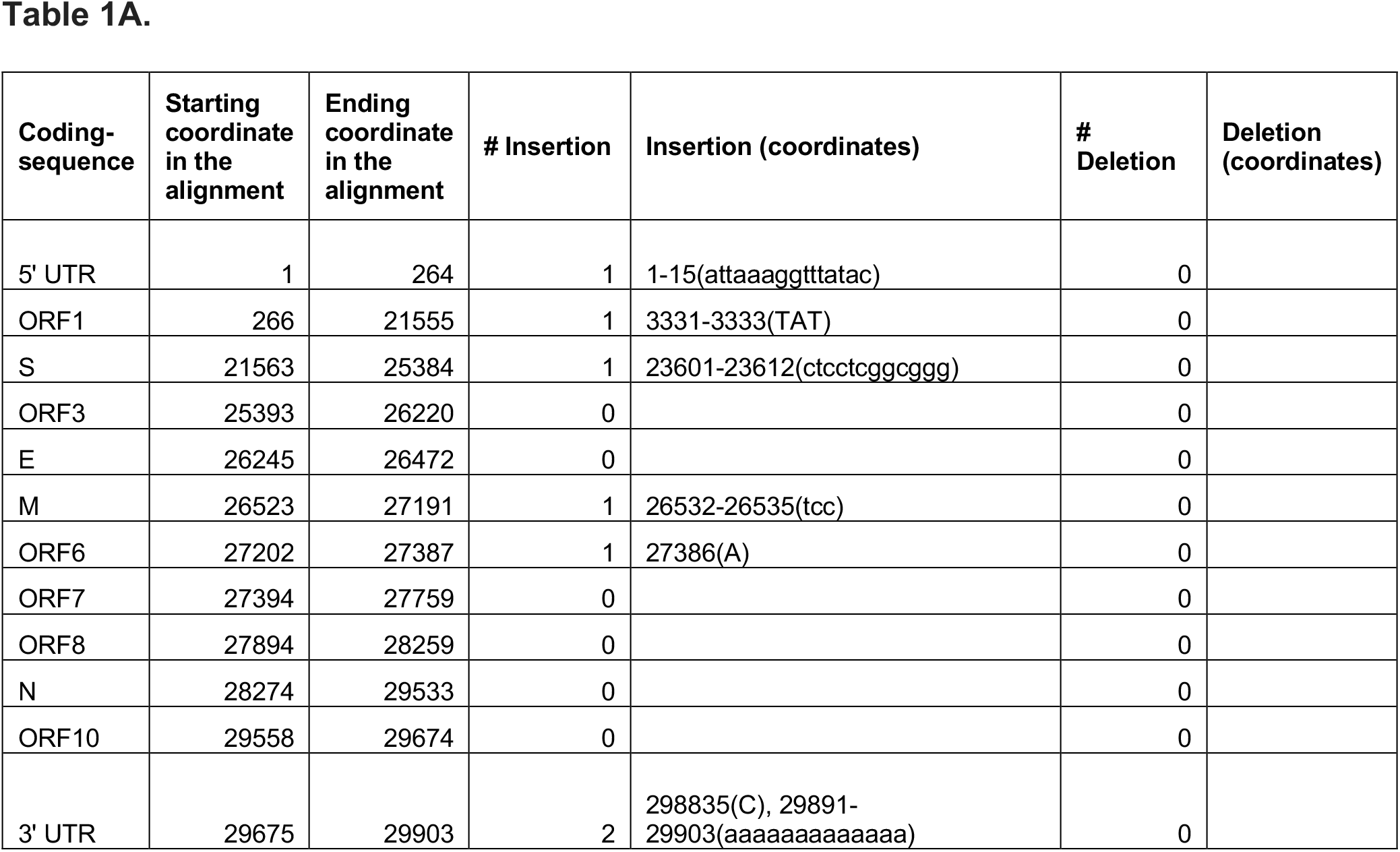
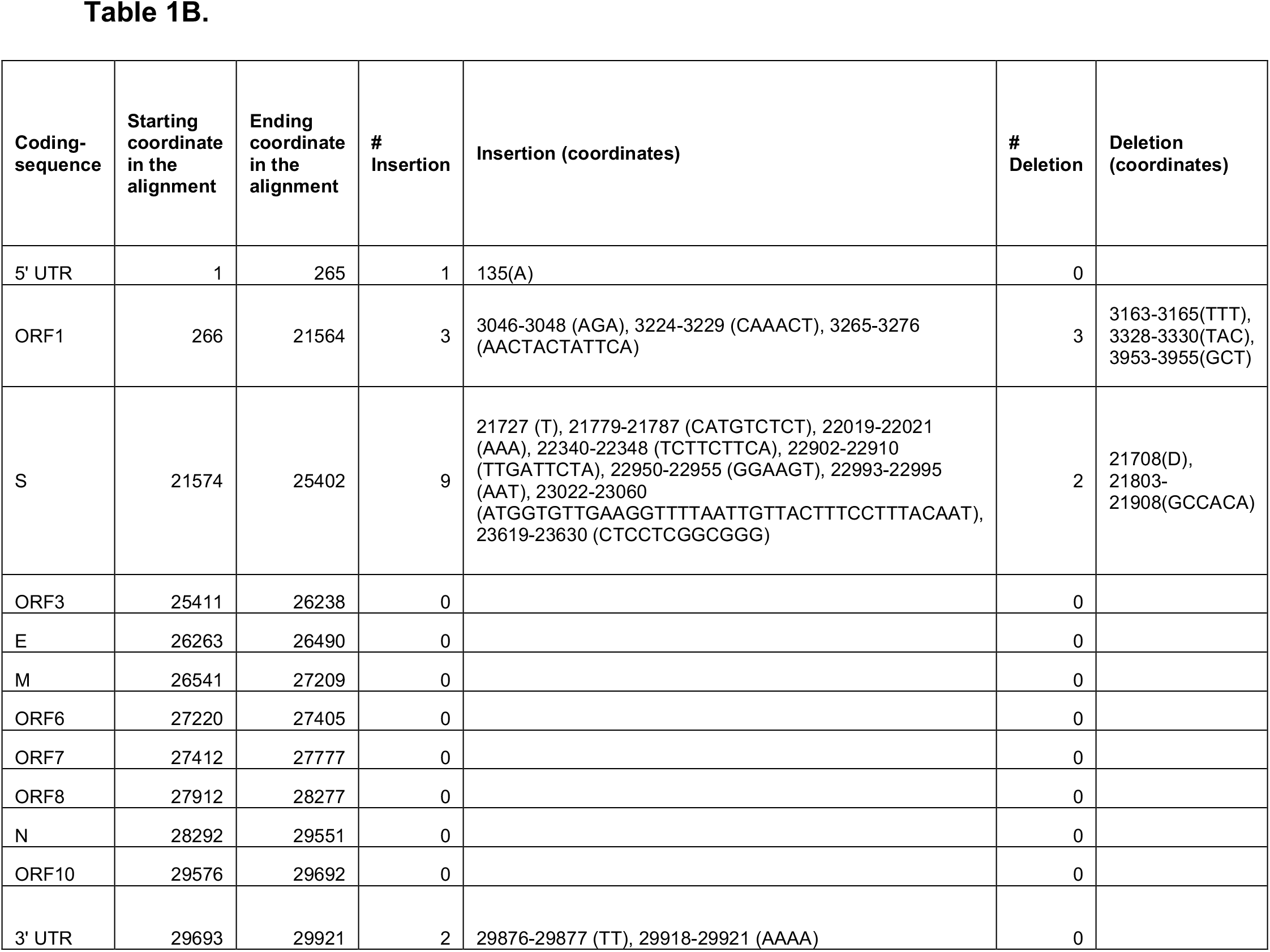
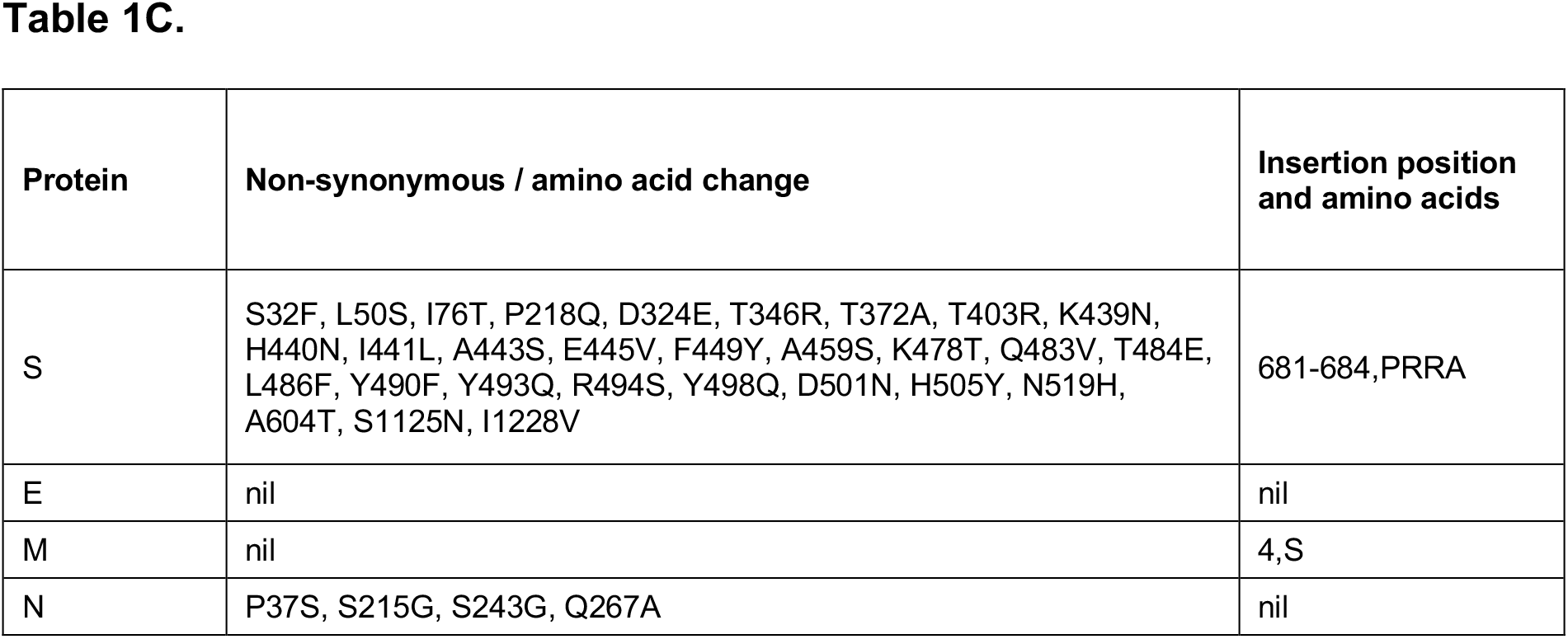
Details of the insertions and deletions (INDELs) in the genome of SARS-CoV-2 (MN908947.3) relative to **A)** bat/Yunnan/RaTG13/2013 (GISAID #EPI_ISL_402131) and **B)** bat-SL-CoVZC45/2017 (MG772933.1) viruses. **C)** List of non-synonymous changes and INDELs in the four structural proteins of SARS-CoV-2 relative to bat/Yunnan/RaTG13/2013.

The genome alignment between the SARS-CoV-2 and bat (bat/Yunnan/RaTG13/2013) coronaviruses showed 2 of 7 insertions (Table 1A) in Spike gene, whereas the alignment of SARS-CoV-2 with distantly related SARS-like bat-SL-CoVZC45/2017 showed 9 of 15 insertions and 2 of 5 deletions (Figure 1C; Table 1B) in Spike gene. Interestingly, the Spike gene has a larger 39 nucleotide insertion (5’-aAT GGT GTT GAA GGT TTT AAT TGT TAC TTT CCT TTA CAA Tca-3’) at genomic location 23022. Our alignment result suggests a possible occurrence of recombination and non-synonymous changes in Spike gene of SARS-CoV-2 which can alter the preference of viral attachment to host receptor. The non-synonymous changes in Nucleocapsid gene likely differentially regulate its binding to virus RNA genome and viral replication. Taken together, our findings implicate that overall genetic differences likely facilitated the novel coronavirus to adapt to humans from bat via altered cell receptor binding, changes in viral genome replication and better evading host immune detection.

### Epitopes in SARS-CoV-2 structural proteins are differentially recognized by HLA-DR alleles

Host immune responses orchestrated by HLA molecules determine the outcome of infectious diseases. Antigen presentation by class II MHC isoforms play a critical role in inducing CD4+ Th1 and Th2 cell-mediated immune responses. It is important to understand the nature of immune selection pressure and evolution of immune escape viral mutants. Since the current outbreak is originated from Wuhan city of China, we set out to study the SARS-CoV-2 epitopes displayed by HLA-DR alleles predominantly present in the ethnic populations of China (PRC), Thailand (TH), Japan (JPN) and Asia-Pacific Region (APR). Thus, we calculated the average binding affinity score of each of the predicted 15-mer peptides from functionally important four structural proteins of SARS-CoV-2.

A total of 134 unique average HBA TCEs were identified in 16 islands/hotspots across the structural protein sequences (Figure 2; Supplemental Table 2). The lowest and highest numbers of average HBA TCEs were identified in nucleocapsid protein (n=11), and spike protein (n=78), respectively. This result reveals that mutations in spike protein may contribute to changes in viral virulence observed in SARS-CoV-2. The overall binding affinity patterns of HLA-DR alleles from different countries were similar, however there are some exceptions: i) no HBA epitope was identified in nucleocapsid protein for TH, JPN and APR dominant HLA-DR allele, while this protein exhibited 3 and 8 TCEs for PRC and Global population set alleles, and ii) HLA-DR alleles for APR and TH recognized lower (n=15) and higher (n=65) number of HBA epitopes, respectively. Nevertheless, delineating common epitopes is crucial for designing universally potent subunit vaccine. Thus, we investigated common epitopes that are recognized by all dominant HLA-DR alleles prevalent in the five ethnic populations. Comprehensively, 8 unique HBA epitopes distributed across Spike (n=2; 232-GINITRFQTLLALHR-246; and 233-INITRFQTLLALHRS-247), Envelope (n=3; 55-SFYVYSRVKNLNSSR-69; 56-FYVYSRVKNLNSSRV-70; and 57-YVYSRVKNLNSSRVP-71) and Membrane (n=3; 97-IASFRLFARTRSMWS-111; 98-ASFRLFARTRSMWSF-112; and 99-SFRLFARTRSMWSFN-113) proteins, were identified (Figure 2; Supplemental Table 2). The high affinity epitopes identified in the functionally important regions of Spike and Membrane proteins could be critical in modulating immune responses to circulating SARS-CoV-2. Many of the low binding affinity regions in the structural proteins may favor viruses to evade host antiviral immunity. Our analysis suggests that a subunit vaccine containing the eight immunodominant HLA-DR epitopes possibly induce effective antiviral T-cell and antibody responses in different ethnic populations.

In summary, the present study provided a detailed genetic analysis of SARS-CoV-2 genome evolution and potential universal epitopes for subunit vaccine development. This SARS-CoV-2 strain might be evolved from the bat-coronavirus by accumulating favorable genetic changes for human infection.

## ACKNOWLEDGEMENTS

We gratefully acknowledge the authors, the originating and submitting Laboratories for their sequence and metadata shared through GISAID and GenBank, on which this research is based.

## SUPPLEMENTAL TABLES

**Supplemental Table 1** - Summary of authors, source laboratories and metadata from GISAID for SARS-CoV-2 genomes analyzed in this study.

**Supplemental Table 2** - Amino acid sequences of high confidence mean HBA (IC <= 10nM) CD4 T-cell epitopes from functionally important regions of the four structural proteins (S, E, M and N) of SARS-CoV-2. The details of protein name, starting and ending positions of epitopes in the given protein, and epitope sequences are provided. The unique epitope sequences for each protein and the universal epitopes commonly recognized by HLA-DR alleles of all five population sets (highlighted in red) are listed at the end of each table.

